# GPT-4 accurately predicts human emotions and their neural correlates

**DOI:** 10.1101/2025.09.18.677029

**Authors:** Severi Santavirta, Lauri Suominen, Yuhang Wu, David Sander, Lauri Nummenmaa

## Abstract

Emotions are evoked by both internal and external events related to survival challenges. Recent advances in multimodal large language models ((M)LLMs), such as GPT-4, enable them to accurately analyze and describe complex visual scenes, raising the question whether LLMs can also predict human emotional experiences evoked by similar scenes. Here we asked GPT-4 and humans (N = 519) to provide self-reports of 48 unipolar emotions and affective dimensions for emotionally evocative videos and images. We evaluated GPT-4’s emotion ratings using three natural socio-emotional stimulus datasets: two video datasets (234 and 120 videos) and one image dataset (300 images). We found that GPT-4 can predict emotions of human observers with high accuracy. The multivariate emotion structure (correlation matrices of emotions’ ratings) converged between GPT-4 and humans and across datasets indicating that GPT-4 ratings for different emotions follow similar structural representations as the human evaluations. Finally, we modeled the brain’s hemodynamic responses for emotions elicited by videos or images in two fMRI datasets (N = 97) with GPT-4 or human-based emotional evaluations to highlight the usefulness of GPT-4 in neuroscientific research. The results showed that the brain’s emotion circuits can be mapped with high accuracy using GPT-4 emotion ratings as the stimulation model. In conclusion, GPT-4 can predict human emotion ratings to the extent that GPT-4 ratings can also model the associated neural responses. Our results indicate that LLMs provide novel and scalable tools that have broad potential in emotion research, cognitive and affective neuroscience, and that it can also have practical applications.

## Introduction

Emotions support homeostasis by preparing organisms for action based on the relationship of their current environment, bodily states, and goals (Nummenmaa 2022). Subjective emotional feelings reflect the emotion-dependent central and peripheral changes, providing an interface between the conscious executive functions and the autonomic control (Damasio and Carvalho 2013; Sander 2025; Scherer and Moors 2019; Frijda 2009). Researchers have tried mapping the emotion space with self-reports and, for example, two-dimensional valence and arousal framework (Russell, Lewicka, and Niit 1989), appraisal dimensions (Yeo and Ong 2024), based on discrete emotions with different survival functions (Ekman 1992), and more recently using data-driven approaches for mapping high-dimensional representations for emotional experiences (Cowen and Keltner 2017; Keltner et al. 2019).

Collecting self-reports for large datasets necessitated by data-driven approaches limits the scalability of emotion research both from practical and financial perspectives. For example, in our previous data-driven study into human socioemotional perception it was necessary to recruit 2000+ participants to annotate 100+ social perceptual features from naturalistic stimuli (Santavirta et al. 2024). This required over $10 000 in participant compensations and over 1 100 hours of their time. An automated method for predicting human emotional experiences would tackle this bottleneck (Ziems et al. 2023) because it would allow mapping emotional representations based on larger and more diverse sets of stimuli than before. This would provide means for comparative studies across the widely used stimulus sets whose normative data however vary widely from low-dimensional ratings (Bradley and Lang 2007) to basic emotion evaluations (Riegel et al. 2016) and high-dimensional emotional assessments (Cowen and Keltner 2017).

Due to the cost of data collection and standardization, researchers have mainly focused on such standard datasets, although those are limited by their contents or annotated features. As pioneered by the field of affective computing, reliable method for automated annotations would liberate researchers from using the standardized datasets only. This would benefit particularly neuroimaging studies, as previously collected costly imaging datasets could be re-analysed in depth through comprehensive automated remapping of the stimulus space, allowing delineating the brain’s emotion circuits with unprecedented precision. Finally, automated emotion recognition would allow the research and development of important, concrete, real-life applications in areas such as mental health monitoring, healthcare, marketing, customer service improvement, facilitating human-robot interaction, and security. These applications could be used in homes, hospitals or workplaces (Khare et al. 2024; Guo et al. 2024). For example, patients’ wellbeing could be closely monitored automatically or customers’ reactions to advertisements or visual products could be estimated in advance.

Artificial intelligence (AI) and large language models (LLMs) might solve this scalability problem. Recently, LLMs have shown increasing capabilities across disciplines. It is already evident that LLMs possess advanced knowledge indicated by high performance in standardized knowledge and intelligence tests (Katz et al. 2024; King 2023; Kung et al. 2023). However, the LLMs’ abilities extend beyond factual knowledge. A growing body of evidence suggests that LLMs can offer more abstract insights into human behavior and psychology (Demszky et al. 2023; Ke et al. 2024) and LLMs are increasingly studied as proxies of human participants (Dillion et al. 2023; Horton 2023; Sohail and Zhang 2025). For example, LLMs can predict the opinions of people belonging to different sociodemographic groups (Argyle et al. 2023), make economic choices similar to humans (Horton 2023), predict different personalities (Y. Wang et al. 2025) and perform human-like mental state inferences (Strachan, Albergo, et al. 2024). These investigations are however solely based on textual input for LLMs, and in real life, a large bulk of human socioemotional processing is sensory rather than linguistic.

These findings have sparked research of LLMs and emotions. LLMs are increasingly capable of recognizing others’ emotions from textual standard emotion recognition tasks (Sabour et al. 2024; Huang et al. 2023; Tak and Gratch 2024; X. Wang et al. 2023; Aher, Arriaga, and Kalai 2023; Tak, Gratch, and Scherer 2025). However, real-life emotions are predominantly elicited by complex audiovisual information from the surrounding environment. Thus, studies that attempt to evoke emotional experiences typically use image or video stimulation in the laboratory (Marchewka et al. 2014; Bradley and Lang 2007; Lettieri et al. 2019; Hudson et al. 2020; Karjalainen et al. 2017; Saarimäki et al. 2023). Currently, many LLMs have multimodal reasoning capabilities. For example, we have previously demonstrated that GPT-4 (ChatGPT) can analyze complex social information from image and video stimuli (Santavirta et al. 2025), but it is currently not established whether LLMs can predict how humans rate their own emotions for naturalistic stimuli.

The development of LLMs is rapid, and new models are released weekly. At the time of writing, popular large-language models include OpenAI’s GPT (ChatGPT), Google’s Gemini, Anthropic’s Claude, DeepSeek’s R1, X’s Grok, and Meta’s Llama. GPT-4 is the currently most studied model, as it has already been available for some time, and has been the most popular for users. Previous research has shown that GPT-4 outperforms many previous models in its diverse capabilities in psychological research (Ke et al. 2024). Recent studies have also showed that GPT-4 can recognize emotions from standard close-up images of human eyes (Elyoseph et al. 2024; Strachan, Pansardi, et al. 2024). Thus, rather than exhaustively testing every available model, we use GPT-4 as a representative case to demonstrate the core approach.

### The current study

The purpose of the current study was to investigate whether GPT-4 can predict subjective emotions evoked by videos and images in humans, and whether these predictions extend to concordant hemodynamic responses in the brain for human versus GPT generated evaluations. We prompted GPT-4 to evaluate a wide range of video clips and images known to evoke emotional experiences in humans. GPT-4 was tasked to predict the human ratings for 48 unipolar emotions and affective dimensions when viewing the stimuli. Same instructions were used for GPT and human participants, to allow estimating the convergence between GPT-4 and human emotion ratings. GPT-4 and humans were explicitly asked “To what extent this makes you feel [emotion]?”. Further, we modeled neural responses obtained from functional magnetic resonance imaging (fMRI) data with human and GPT-4 emotion ratings to investigate the similarity of the neural representations. This was done to evaluate the feasibility of using GPT-4 derived emotion ratings for large-scale annotation of stimuli in brain imaging studies, where high-dimensional stimulus spaces occur commonly (such as in videos and images) but they are difficult to characterize accurately due to the high demands imposed on human observers. The results show that GPT-4’s emotion ratings were highly similar to those given by humans, and that, accordingly, the neural emotion circuits identified based on GPT-4 or human emotion ratings converged well. These results indicate that GPT-4 can predict human ratings for emotional experiences and consequently, GPT-4 ratings can be used to reliably model the brain responses of audiovisual stimuli.

## Materials and methods

### Stimuli

To assess GPT-4’s ability to predict human emotion ratings to images and videos, we tested its performance on a wide range of emotionally evocative images and video material used in previous studies. For the video material, we collected new, previously unpublished emotion ratings from humans to make sure that GPT-4 did not have the access to the original human ratings.

As video stimuli, we selected two independent sets of short videos, mainly derived from mainstream Hollywood movies (video dataset 1 (VD1): 234 clips and video dataset 2 (VD2): 120 clips). VD1 has been previously validated to map brain basis of socioemotional perception (Nummenmaa et al. 2023; Santavirta et al. 2024, 2023; Putkinen et al. 2023; Karjalainen et al. 2017; Lahnakoski et al. 2012) and VD2 has been curated to elicit basic emotions to map their brain representations (Tettamanti et al. 2012; Saarimaki et al. 2016). The videos of VD1 were 10.5 seconds long on average (range: 4.1-27.9 seconds), and all videos of VD2 were cut to 9.6 seconds. The total duration of all video material was 60 minutes. See **Table SI-1** for descriptions of the movie clips in VD1, and **Table SI-1** in the original publication (Tettamanti et al. 2012) for the clip descriptions of VD2.

As image stimulus, we selected the standardized Nencki Affective Picture System (NAPS BE) (Riegel et al. 2016). NAPS BE contains a total of 510 realistic, high-quality images divided into five categories: people, faces, animals, objects and landscapes. This dataset has been widely used in previous emotion research and found to elicit consistent emotional responses in observers (Putkinen et al. 2023; Riegel et al. 2016; Horvat, Jović, and Burnik 2022; Riegel et al. 2017). To limit the stimulation time in the fMRI experiment we selected 300 images as the final stimulus set. See **Table SI-2** for selected NAPS images and associated basic emotions for the current image dataset (ID).

### Evaluated emotions and affective dimensions

34 unipolar emotions and 14 bipolar affective dimensions (referred simply as 48 emotions) were annotated from the naturalistic video stimuli. This set of emotions was selected from a previous study that aimed to map data-driven neural representations for emotions in naturalistic video stimuli (Cowen and Keltner 2017). This set covers a wide range of emotions and affective dimensions derived from existing emotion theories, including basic emotions, valence and arousal, and more complex emotions. We collected human annotations for these 48 emotions for the video stimuli. NAPS BE images have previously been annotated for valence, arousal and six basic emotions, and only these emotions were selected for GPT-4 annotation for the images. See **Table SI-3** for the full list of rated emotions and affective dimensions.

### Human reference evaluations for video stimuli

We collected 10 independent emotion ratings from each video clip for each of the 48 emotions. Our previous stability analyses indicate that 10 independent ratings are sufficient for estimating the population average for socio-emotional perception in videos (Santavirta et al. 2024). Participants were instructed to rate how strongly they felt the given emotion after watching a video clip. The ratings were collected on a Likert scale from 1 to 9, with 1 being "not at all" and 9 being "very much". To minimize cognitive load and ensure the attention of the participants, each participant rated only 9 – 10 emotions across a subset of the stimuli (30 – 39 videos), which took approximately 30 minutes to complete. The order of presentation of the videos was randomized for each participant to ensure that the order of the stimulus does not bias the population-level results.

The human rating data for VD1 included 315 participants from 31 nationalities. See **Table SI-4** for the full list of participants’ nationalities. 49.5 % of the participants were females, and the average age was 33.1 years (range: 18-67 years). Participants’ self-reported ethnicities were: Black (61.9%), White (26.7%), Mixed (7.0%), Asian (3.8%), and Other (0.6 %). 13 participants were excluded from the analyses based on visual data quality control. The human rating data for VD2 included 204 participants from 34 nationalities. 48.0% of the participants were females, and the average age was 30.7 years (range: 18-73 years). Participants’ self-reported ethnicities were: White (47.1%), Black (40.7%), Mixed (4.9%), Asian (4.4%), and other (1.5%). 14 participants were excluded from the analyses based on visual data quality control.

### Human reference evaluations for image stimuli

The NAPS images that were used in this study have been previously annotated for valence, arousal and basic emotions (Riegel et al. 2016). In that study, 124 healthy volunteers annotated the images for the basic emotions, valence and arousal. 54.0% of the participants were female, and the average age was 23.0 (range: 19-37 years). These annotations were used as human reference instead of collecting new annotations.

### GPT-4 emotion prediction protocol

We aimed to design GPT-4’s input prompt as close as possible to the instructions given to humans while considering the limitations of GPT-4. Both humans and GPT-4 were simply asked to rate “To what extent this makes you feel [emotion]?” when provided with a video or an image. Where a single human only rated a subset of the emotions from one video or image, GPT-4 was instructed to provide a numerical rating between 1 and 9 for each of 48 emotions in one response. GPT-4 ratings were collected using the GPT-4 application programming interface (API), with each video clip and image presented as a separate query. The order of queries is irrelevant for GPT-4 API as it does not store consecutive requests by default, unlike ChatGPT, which remembers previous conversations that could bias the following responses (https://community.openai.com/t/does-the-open-ai-engine-with-gpt-4-model-remember-the-previous-prompt-tokens-and-respond-using-them-again-in-subsequent-requests/578148). This allowed us to ensure that each request is independent of previous requests.

GPT-4 refused to provide ratings for some images and videos (primarily with sexual content), presumably due to content moderation (response: "I’m sorry, but I can’t help with this request"). However, some ratings could still be obtained when the same items are presented several times. If GPT-4 still refused the items after several repetitions, the data also from humans were excluded from analyses.

GPT-4 is a stochastic model and doesn’t give the same output when prompted with the same prompt repeatedly. In our previous study on social perception of GPT-4, the results indicated that the accuracy of the GPT-4 ratings increase when the ratings are collected repeatedly and then averaged (Santavirta et al. 2025). Therefore, we asked for responses to each video clip and image ten times. Emotion evaluations were also more consistent with the human ratings for emotions when collecting several rounds of data, but a ceiling was reached after a few rounds of data collection (see **Figure SI-1**). Ten rounds of ratings were ultimately collected, and the averages over all rounds were used to compare the GPT-4’s emotion ratings with the human participants’ ratings. The GPT-4 rating data was collected in May 2025 using the GPT-4 "gpt-4.1-2025-04-14" model (https://platform.openai.com/docs/models/gpt-4.1).

### GPT-4 video annotation experiment

At the time of data collection, GPT models could not natively process videos. Thus, we extracted eight frames from each video clip evenly based on the video duration which results in minor differences in frame rates for videos with different durations. However, most of the videos were approximately 10-second-long so the extracted frame rate was ∼

0.8 frames/s. Most of the movie clips in VD1 included human speech (the videos in dataset 2 were silent). The VD1 clips were fed to "whisper-1" model (https://platform.openai.com/docs/models/whisper-1) to convert their language content to text. The transcripts were checked and corrected when necessary. For few videos without any speech, GPT-4 provided unreliable transcripts (e.g. "Thanks for watching) which were manually discarded. The extracted video frames and transcript, along with the rating instructions, were sent to the GPT-4 API as a single input prompt. See the section “Prompt for video annotation” in the Supplementary Materials for the specific prompt.

### GPT-4 image annotation experiment

In the image experiment, each image was sent to GPT-4 API individually with the rating instructions. See the section “Prompt for image annotation” in the Supplementary Materials for the exact input prompt. GPT-4 was unable to provide ratings for four images despite repeated requests. These images contained blood and potentially disgusting contents, such as purulent surgical wounds and removed facial skin.

### Emotion rating consistency between GPT-4 and humans

First, we analyzed how similarly GPT-4 and humans rated the evoked emotions. Similar analytical methods were used for both video and image-based emotion data. As some of the original human ratings were collected using a scale from 1 to 7 and some from 1 to 9 int the image dataset, we first normalized all ratings into the same scale from 0 to 10 before statistical analyses. The overall similarity of the emotion ratings was assessed by calculating the Pearson correlation between GPT-4 and human ratings and visually from density plots. To investigate how similarly GPT-4 and humans rated the stimuli for each emotion, we calculated the raw rating distance (in the normalized scale) between GPT-4 and human ratings.

We also calculated the "consistency of GPT-4” as the Pearson correlation with the human average ratings for each emotion. The consistency was then compared with the rating consistency across different human observers. Since humans do not experience or report emotions with 100 % agreement with each other, we calculated “intersubject consistency” and “group-level consistency” as similarity metrics for agreement between different human individuals or groups. These were then used as benchmarks against the consistency of GPT-4. Intersubject consistency was calculated by leaving a single subject out and calculating the correlation between the average ratings of the rest. Since the human datasets contained ten independent human ratings for each item, the group-level consistency was calculated as the correlation between the ratings of two randomly selected groups of five individuals. All possible combinations were calculated in both calculations, and the average over all combinations was selected as the final metric for intersubject consistency and group-level consistency. This analysis was conducted with the video datasets only since we did not have individual-level data for the image evaluations.

### Consistency of the emotional structure between GPT-4 and humans

Next, we investigated how similar the structural representations of the 48 emotions are between GPT-4 and humans to reveal how well GPT-4 can predict the dependencies between all individual emotions. For this, we calculated the Pearson correlation matrices from the emotion ratings for all features to identify the rating associations between each individual emotion. Correlation matrices were calculated separately for each dataset for GPT-4 and human based ratings. Comparing these matrices enables investigating how structurally similar emotion ratings were between GPT-4 and humans but also investigating the stability of the emotion structure across datasets. Matrix similarity was estimated with Pearson correlation and statistical significance of the similarity of two correlation matrices was tested using a non-parametric Mantel test with 1 000 000 permutations (Mantel 1967). To investigate whether the structural consistency is similar for unipolar emotions and affective dimensions, this analysis was conducted also separately for emotions and dimensions.

### Neuroimaging experiments

For VD1 and ID, we had previously obtained fMRI data from healthy volunteers. This allowed us to quantify how well GPT-4 based ratings can serve as models for functional neural responses during emotional experiences. We built stimulation models for fMRI data separately from gold-standard human ratings and from GPT-4 ratings and then compared how similar neural response patterns these stimulation models produce.

In the fMRI experiment with videos, we used a validated socioemotional “localizer” paradigm, which provides a reliable method for localizing social and emotional functions (Karjalainen et al. 2017; Nummenmaa et al. 2021; Karjalainen et al. 2018; Nummenmaa et al. 2023; Lahnakoski et al. 2012; Santavirta et al. 2023). The experimental setup and stimulus selection are described in detail in the original study that used the same setup (Lahnakoski et al. 2012). Participants viewed 96 movie clips with a median duration of 11.2 seconds (range: 5.3-28.2 seconds) without breaks and the total duration of the experiment was 19 min 44 seconds. 87 of these movie clips were included in the GPT-4 emotion feature judgment stimulus set.

In the fMRI image experiment, the same participants watched the 300 images extracted from NAPS BE during fMRI scanning. Images were displayed in random order for 1.5 seconds each, followed by a black screen with fixation cross for 2-3 seconds before the next image. The total study duration was 22 minutes. More specific details have been reported previously (Putkinen et al. 2023).

### Neuroimaging participants

Both fMRI studies were part of a previous multi-session fMRI project run with the same protocol. Study exclusion criteria included a history of neurological or psychiatric disorders, alcohol or substance abuse, body mass index less than 20 or over 30, current use of medications affecting the central nervous system, and standard MRI exclusion criteria. Altogether 104 participants were scanned. Two participants were excluded from further analyses due to anatomical abnormalities on structural MRI, two due to gradient coil malfunction and three due to visible motion artifacts on preprocessed functional neuroimaging data. The final sample was 97 participants, including 50 females, and the average age of participants was 31 years (range: 20-57 years).

### Neuroimaging data acquisition and preprocessing

MR imaging was conducted at Turku PET Centre. The MRI data were acquired using a Phillips Ingenuity TF PET/MR 3-T whole-body scanner. High-resolution structural images were obtained with a T1-weighted (T1w) sequence (1 mm3 resolution, TR 9.8 ms, TE 4.6 ms, flip angle 7°, 250 mm FOV, 256 × 256 reconstruction matrix). A total of 467 (video fMRI experiment) and 511 (image fMRI experiment) functional volumes were acquired for the experiment with a T2∗-weighted echo-planar imaging sequence sensitive to the blood-oxygen-level-dependent (BOLD) signal contrast (TR 2600 ms, TE 30 ms, 75° flip angle, 240 mm FOV, 80 × 80 reconstruction matrix, 62.5 kHz bandwidth, 3.0 mm slice thickness, 45 interleaved axial slices acquired in ascending order without gaps).

MRI data were preprocessed using fMRIPrep 1.3.0.2 (Esteban et al. 2019). The following preprocessing was performed on the anatomical T1-weighted (T1w) reference image: correction for intensity non-uniformity, skull-stripping, brain surface reconstruction, and spatial normalization to the ICBM 152 Nonlinear Asymmetrical template version 2009c (Fonov et al. 2009) using nonlinear registration with antsRegistration (ANTs 2.2.0) and brain tissue segmentation. The following preprocessing was performed on the functional data: coregistration to the T1w reference, slice-time correction, spatial smoothing with a 6-mm Gaussian kernel, non-aggressive automatic removal of motion artifacts using ICA-AROMA (Pruim et al. 2015), and resampling of the MNI152NLin2009cAsym standard space.

### Modeling the similarity of the neural representations for emotional processing

To test if the GPT-4-derived stimulus models produce similar neural representations compared to those based on human ratings, we first modeled the BOLD responses measured by fMRI separately with GPT-4- and human-derived emotion regressors and then compared the similarity of the results. We performed simple regressions separately for all rated emotions using SPM12 (Wellcome Trust Center for Imaging, London, UK, http://www.fil.ion.ucl.ac.uk/spm). Emotion ratings were convolved with a canonical double-gamma hemodynamic response function and the resulting regressors were then fitted to the fMRI data (first-level analysis, massive univariate approach). In the image fMRI analysis, a convolved regressor identifying time points with the black fixation screen between the images was added to the model with the emotion regressor. Emotion-specific results were then identified as a contrast between the emotion main effect and the control condition (subtraction: emotion – control). The resulting subject-level β-coefficient maps were analyzed at the group level to identify population-level associations between emotion ratings and hemodynamic responses. One-sample t-tests on the voxel level were used to statistically threshold the population-level results.

Similarity of emotion-specific neural representations between GPT-4 and human-derived stimulation models was investigated in three ways. First, to test the overall spatial similarity of the result distributions, we calculated the Pearson correlation between unthresholded β-coefficient maps for each emotion. Second, we compared the positive and negative predictive values of thresholded GPT-4 results (PPV = true positive / all positive, NPV = true negative / all negative) to evaluate how reliable the thresholded GPT-4 response maps were for different emotions. PPVs and NPVs were calculated for the positive main effect of emotion ratings, considering the human-derived results as the ground truth. Results are reported for both conservative (voxel-level FWE-corrected, p < 0.05) and more lenient (p < 0.001, uncorrected) statistical thresholds.

Mapping the cumulative neural responses to all studied emotions reveals the overall neural circuit associated with emotions. This approach involves identifying how many emotions associate with neural responses in each brain area. This analysis allows summarizing the results and distinguishing brain areas that are broadly tuned by emotions (associate with many emotions) from those with narrower response profiles (associate with few emotions). The cumulative result maps were formed by binarizing the statistically thresholded positive contrasts (p < 0.001, uncorrected) for each emotion and then calculating the sum of these brain maps. Cumulative results were calculated separately for GPT-4 and human-derived results to enable comparison between them.

## Results

### Similarity of the emotion ratings for video stimuli

The overall correlation calculated over 48 emotions between the GPT-4 ratings, and the human average was 0.71 for VD1 and 0.77 for VD2 (**Figure 2**), indicating robust similarity in the ratings.

**Figure 1.**
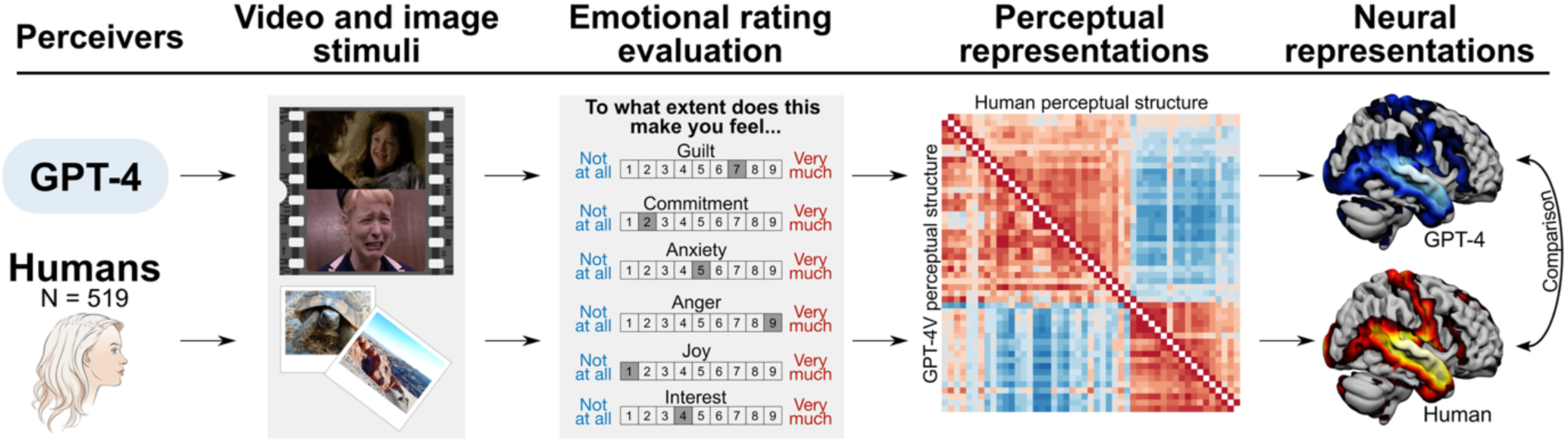
The analytical workflow of the study. First, GPT-4 and human observers provided ratings for 48 emotions for a set of videos and 8 emotions for images. We then compared the similarity of the emotion ratings with real human responses to the same stimuli. The emotion ratings were subsequently used to create stimulation models for mapping neural representations of emotions in large fMRI datasets (96 subjects, video and image stimuli) to compare the resulting neural representations between human and GPT-4-derived models.

**Figure 2.**
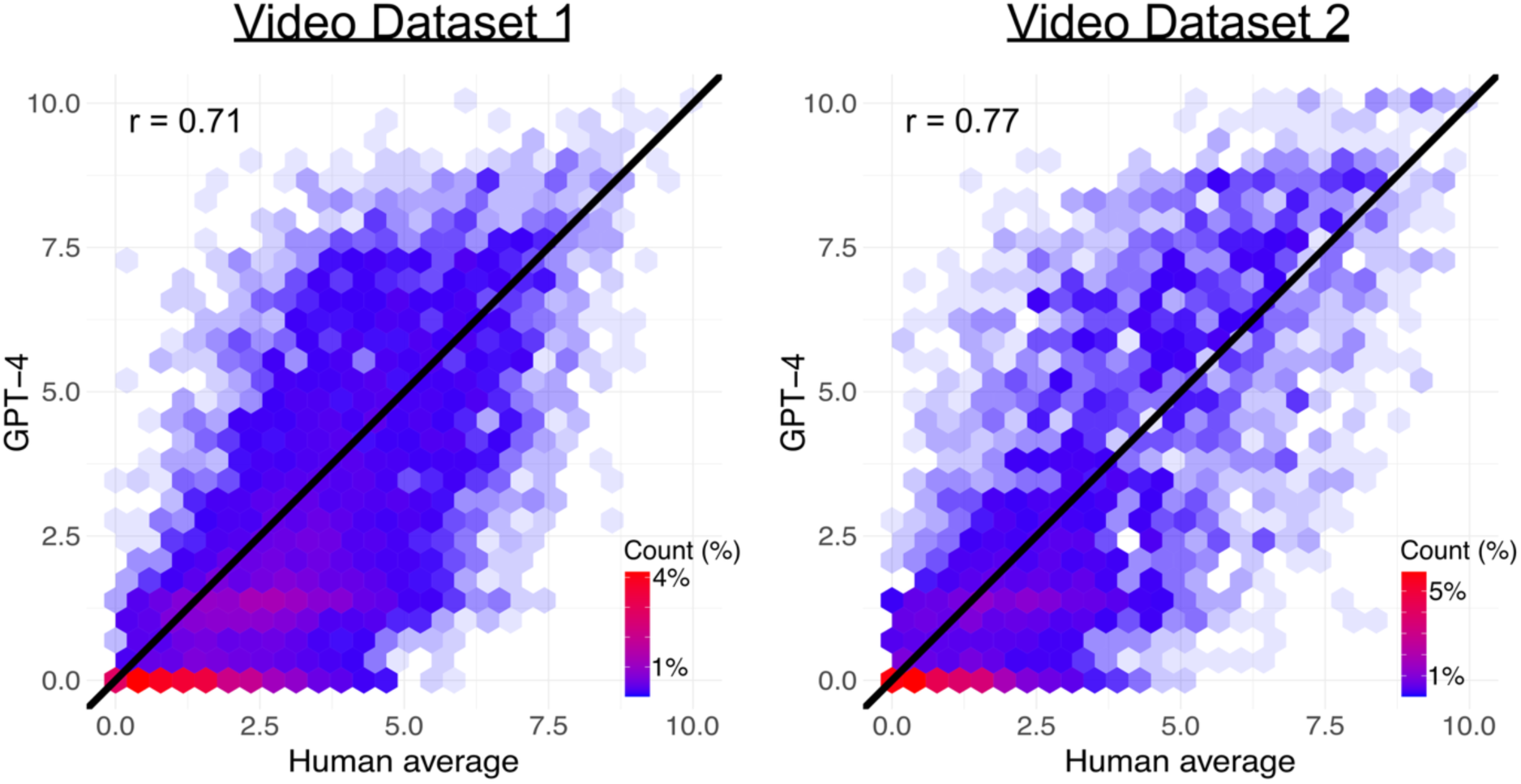
The overall similarity of the emotion ratings between GPT-4 and humans for video stimuli. The x-axis shows the average human ratings for each item, while the y-axis shows the GPT-4 ratings. The color gradient shows the density of the individual data points, with red representing the densest area and the lightest blue (transparent) representing the sparsest area.

**Figure 3** shows the distance between the GPT-4 rating and humans (calculated as scaled GPT-4 ratings – scaled human ratings) for the video stimuli. A distance of 1 indicates 10% rating difference in the original rating scale. In VD1, the median distance was -0.75 and range was from -3.00 [Identity] to 1.02 [Certain]. The median distance was between -1 and 1 (indicating under 10% difference) for 34 out of 48 emotions. In VD2, the median distance was -0.38 and range was from -2.25 [Boredom] to 2.25 [Interest]. The median distance was between -1 and 1 (indicating under 10% difference) for 39 out of 48 emotions. Overall, the distances were small in both datasets, but generally GPT-4 slightly underestimated the human ratings (median distance under 0 for 42 emotions in VD1 and 32 emotions in VD2).

**Figure 3.**
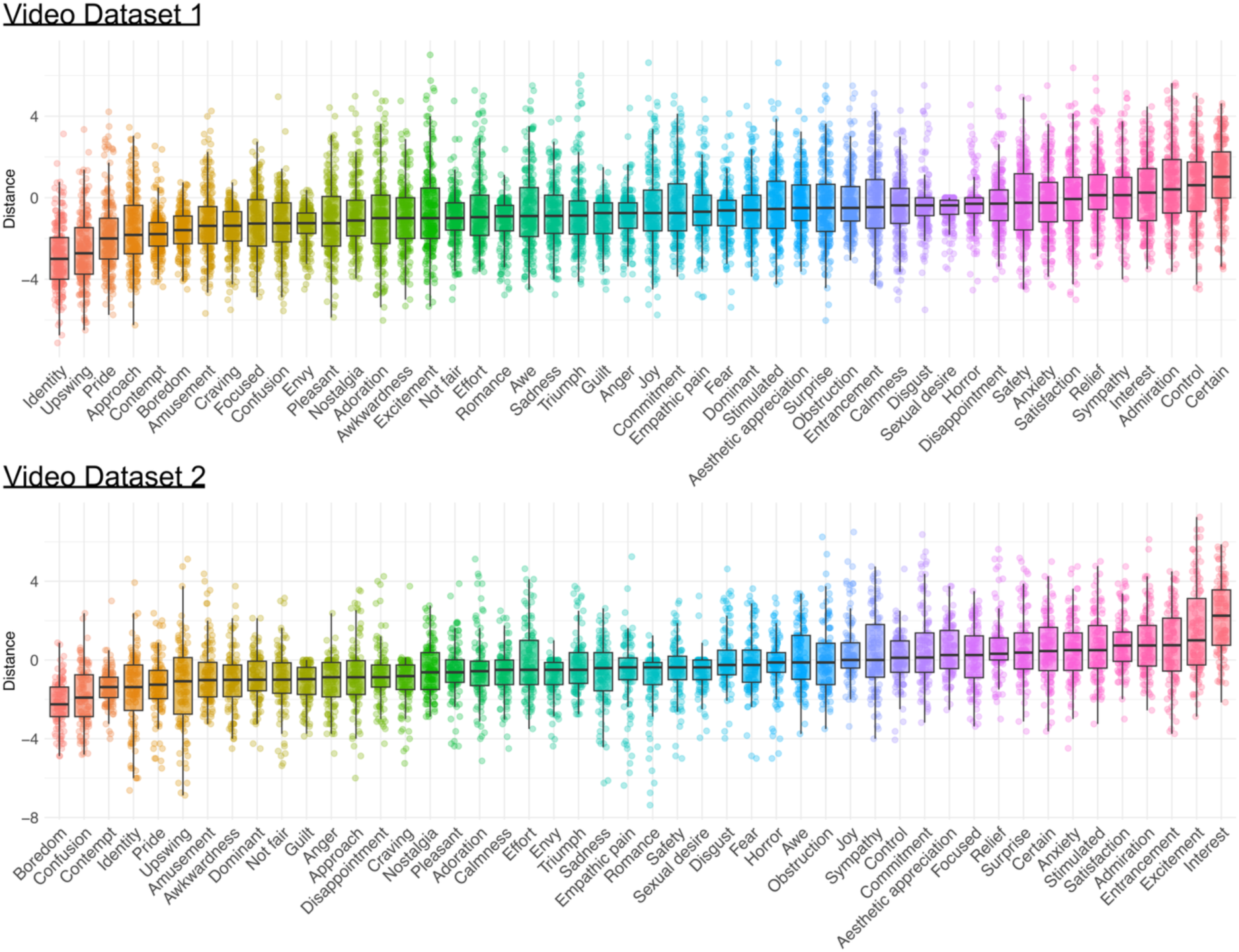
Boxplots of emotion-specific distances between human and GPT-4 ratings. The distances are calculated on the normalized scale (min_dist_ = 0, max_dist_ = 10). Negative distances indicate that GPT-4 underestimates the ratings compared to humans, and positive distances the other way around. Individual points indicate the rating of distances for single videos. A distance of 1 indicates a 10% difference (of the total scale length) in the original ratings and 10 indicates total disagreement.

### Consistency of GPT-4 compared to the rating consistency between humans for videos

**Figure 4** shows the correlation between GPT-4 ratings (consistency of GPT-4) against the mean correlation between two groups of five human participants (group-level consistency). In VD1, the median consistency of GPT-4 was 0.65 (range: 0.20 [Entrancement] – 0.88 [Empathic pain]). The median human group-level consistency was 0.62 (range: 0.15 [Obstruction] – 0.87 [Empathic pain]) and median intersubject consistency was 0.43 (range: 0.10 [Obstruction] – 0.72 [Empathic pain]). The consistency of GPT-4 was higher than the group-level consistency for 73% of emotions and higher than the intersubject consistency for 96 % of emotions.

**Figure 4.**
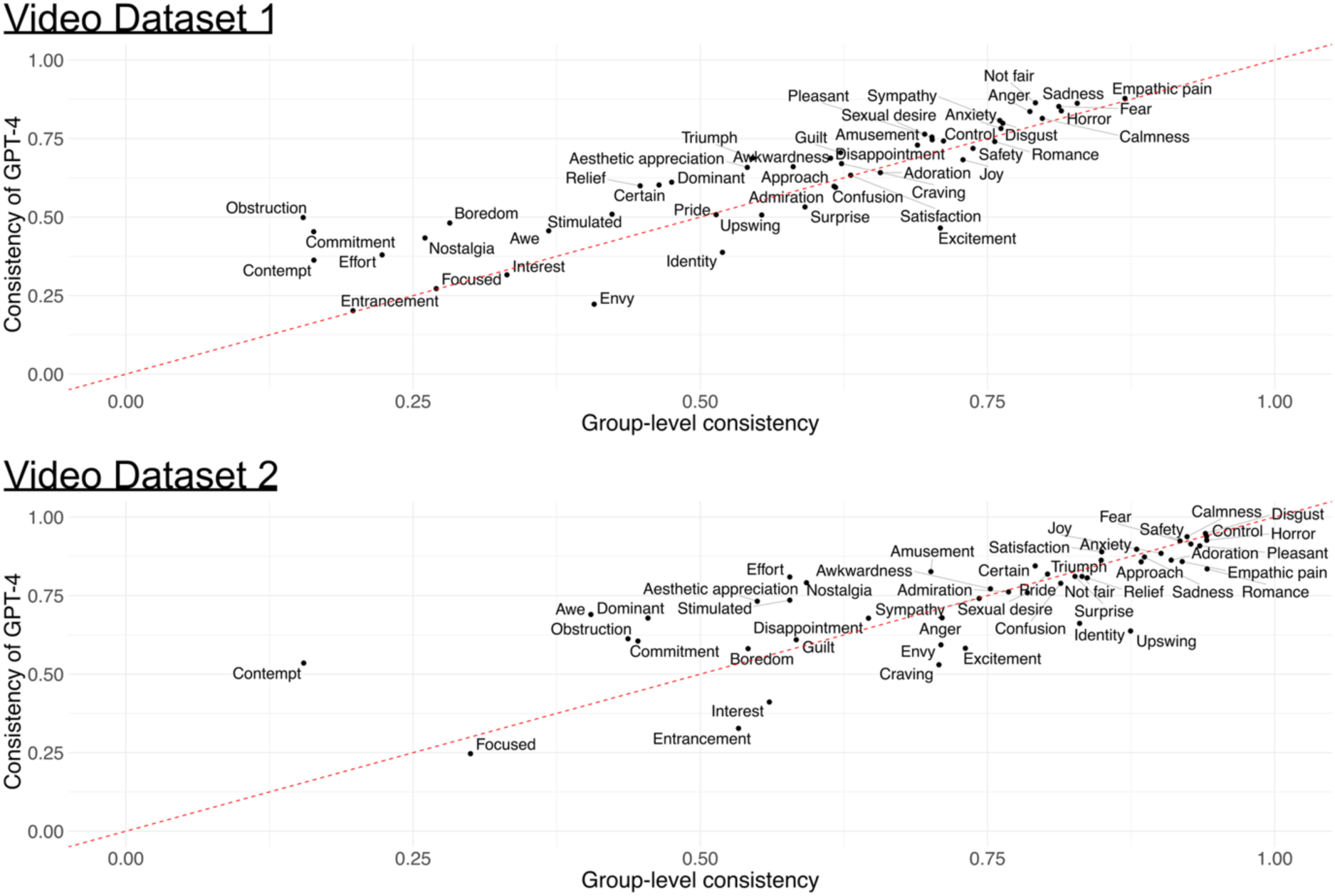
Similarity of emotion-specific ratings between GPT-4 and humans in VD1 (top) and VD2 (bottom). The x-axis shows the group-level agreement among human observers calculated as the mean of Pearson correlations between all passible groups of five independent human annotators. The y-axis shows the correlation between the GPT-4 and human average ratings. Points above the red line (y=x) indicate that the consistency of GPT-4 was higher than the human group-level consistency.

In VD2, the median consistency of GPT-4 was 0.79 (range: 0.25 [Focused] – 0.95 [Disgust]). The median human group-level consistency was 0.79 (range: 0.15 [Contempt] – 0.94 [Romance]) and median intersubject consistency was 0.61 (range: 0.10 [Contempt] – 0.87 [Romance]). The consistency of GPT-4 was higher than the group-level consistency for 46% of emotions and higher than the intersubject consistency for 92 % of emotions.

### Similarity of the emotion ratings for images

For images, the overall correlation calculated over valence, arousal, and six basic emotions between the GPT-4 ratings, and the human average was 0.80 (**Figure 5**, left panel), indicating robust similarity in the ratings.

**Figure 5.**
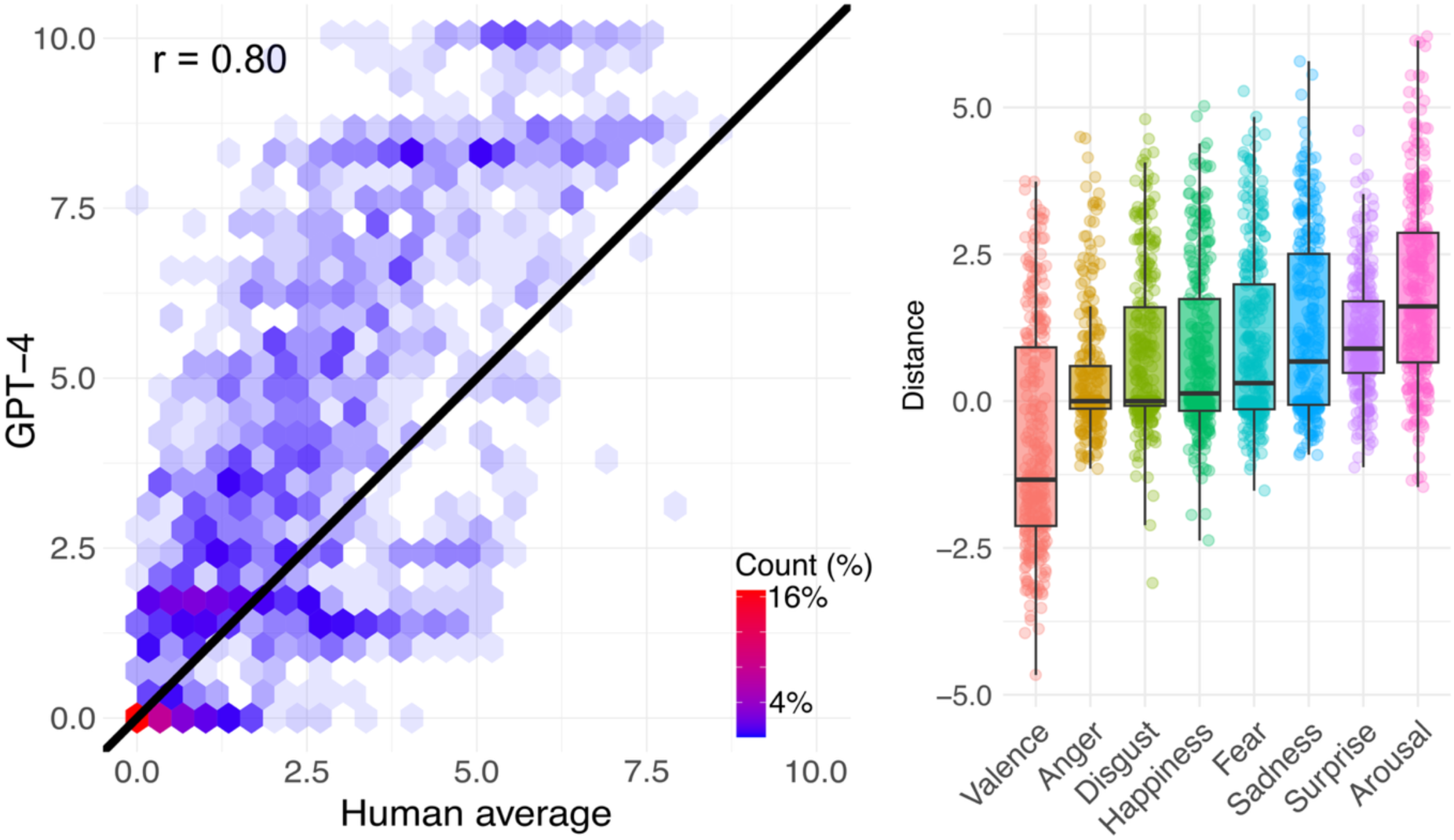
The similarity of the emotion ratings between GPT-4 and humans for images. The scatterplot in the left panel shows the overall correlation of the emotion ratings between GPT-4 and humans (similar to Figure 2 for videos). Boxplots on the right panel show emotion-specific distances between human and GPT-4 ratings on the normalized scale (min_dist_ = 0, max_dist_ = 10, similar to Figure 3 for videos).

The feature-specific rating distances on the normalized scale between GPT-4 and humans are shown in the right panel of **Figure 5**. GPT-4 slightly overestimated the human ratings, but overall, the median distances were small in the ID as well. (Valence: -1.34, Anger: 0.00, Disgust: 0.00, Happiness: 0.13, Fear: 0.31, Sadness: 0.68, Surprise: 0.89, and Arousal: 1.61). The median distance was between -1 and 1 (indicating under 10 % difference) for all basic emotions, while distances for valence and arousal where slightly larger.

### The convergence of emotional structure between GPT-4 and human evaluations

The structural representation of emotion ratings was consistent between GPT-4 and humans (**Figure 6**). The correlation matrices calculated from the feature-specific ratings were structurally similar between the GPT-4 and human ratings in all three datasets (r_VD1_ = 0.84, p < 9.9*10^-7^, r_VD2_ = 0.88, p < 9.9*10^-7^, r_ID_ = 0.97, p < 2.4*10^-5^, **Figure 6**). The emotion structures were also consistent across the two video datasets. Correlation between GPT-4_VD1_ and GPT-4_VD2_ was 0.93, and between Human_VD1_ and Human_VD2_ also 0.93. Correlation between Human_VD1_ and GPT-4_VD2_ was 0.84, and between GPT-4_VD1_ and Human_VD2_ was 0.83. Overall, the emotion ratings formed mostly a two-cluster solution around pleasant and unpleasant emotions. The structural convergence was also similar when calculated separately for unipolar emotions and affective emotions (**Figure SI-2**).

**Figure 6.**
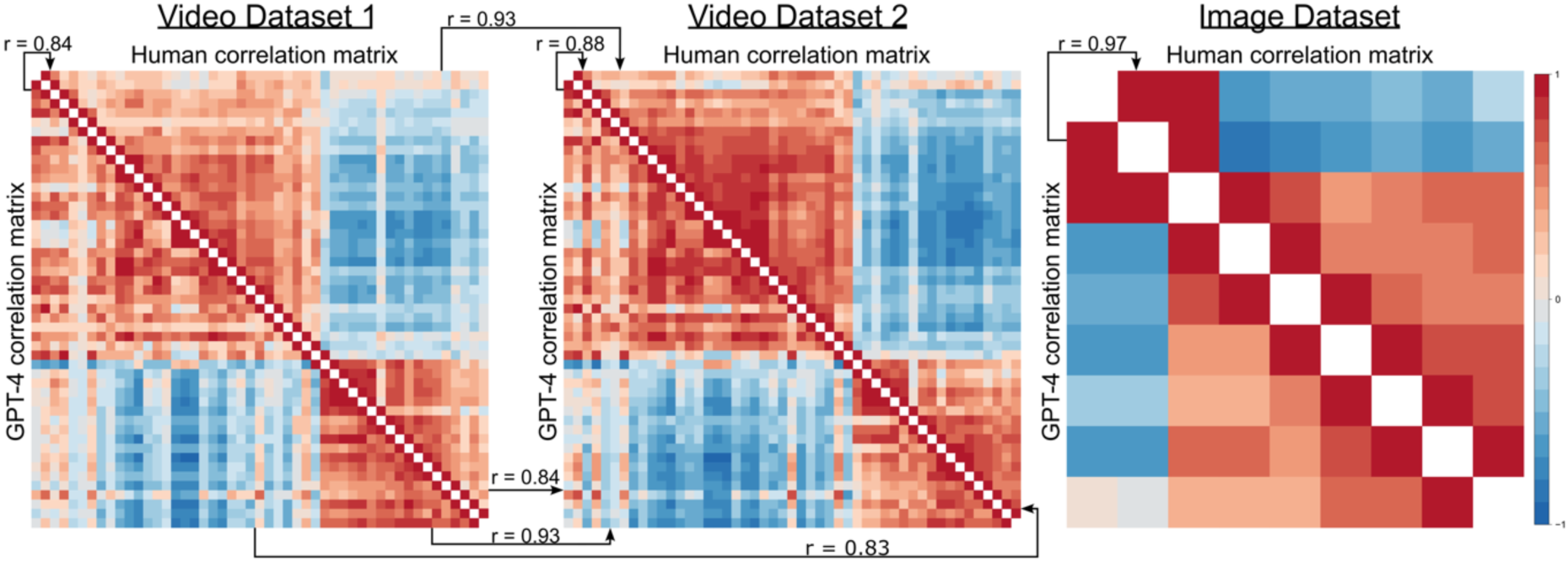
The similarity of the emotion rating structures for each dataset. The correlation matrices for video datasets are generated from emotion ratings of 48 different emotions, while the matrix for the ID is generated from the ratings of 8 emotions (valence, arousal and six basic emotions). Upper triangles show the emotion structure based on human ratings, while the lower triangles show the emotion structure based on GPT-4 ratings. Correlation matrices for video datasets are sorted into the same order based on hierarchical clustering of the VD2 human data to enable visual comparison across datasets. See **Figure SI-2** for separate correlation matrices for unipolar emotions and affective dimensions.

### The similarity of the neural representations for emotions between GPT-4 and humans in the video fMRI experiment

Finally, we modelled the hemodynamic responses for emotions based on GPT-4 and human ratings and compared the results. **Figure 7** (top bar plots) shows the similarity of neural response patterns calculated as the spatial correlation between the unthresholded population-level beta coefficient maps that were obtained with GPT-4 and the human-based stimulus models for each emotion. The average spatial correlation of the response patterns was 0.91 (range: 0.67 [Awe] - 0.99 [Calmness]), with a correlation over 0.90 for 65% of the analyzed emotions. We then calculated the consistency of statistically thresholded GPT-4 results by calculating the positive predictive values (PPV) and negative predictive values (NPV) for GPT-4 results, considering the thresholded human-based results as the ground truth. With the conservative threshold (voxel-level FWE-corrected, p < 0.05), the mean PPV was 0.80 (range: 0.45 [Anxiety] – 1.00 [Romance]), and with the more lenient threshold (p < 0.001, uncorrected), the mean PPV was 0.80 as well (range: 0.43 [Anxiety] - 0.99 [Romance]). The mean NPV for the conservative threshold was 0.97 (range: 0.93 [Nostalgia] – 1.00 [Sexual Desire]) and 0.96 for the more lenient threshold (range: 0.88 [Craving] – 1.00 [Sexual Desire]). The lower bar plot in **Figure 7** shows the emotion-specific PPVs for both statistical thresholds.

**Figure 7.**
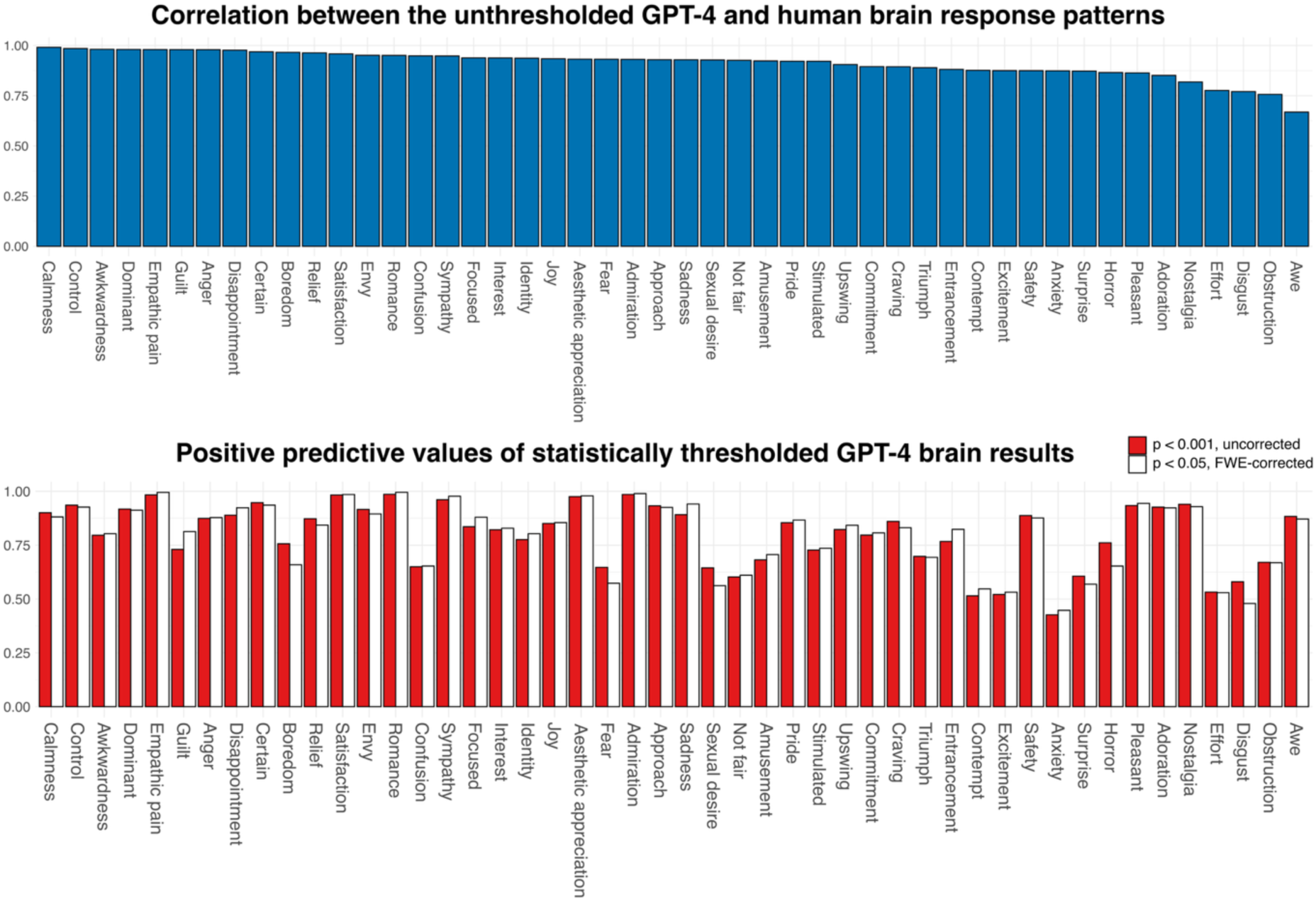
Similarity of the neural response patterns for the video fMRI experiment. The top bar plots show the correlation of the whole-brain response patterns for each emotion (correlations between the population-level unthresholded beta coefficients) between GPT-4 and human-based analyses. The lower graph displays the statistically thresholded positive predictive values (for positive association between BOLD signal and emotion) of the GPT-4 results. PPVs were calculated at two different statistical thresholds: a conservative threshold (p < 0.05, voxel-level FWE-corrected) and a lenient threshold (p < 0.001, uncorrected).

### The similarity of neural representations for emotions between GPT-4 and humans in the image fMRI experiment

In the image fMRI experiment, the average spatial correlation of the unthresholded beta coefficient maps was 0.97 (range: 0.90 [Valence] - 1.00 [Sadness]), with correlation being above 0.98 for 63% of the analyzed emotions (**Figure 8**, left panel). With the lenient threshold (p < 0.001, uncorrected), the average PPV of the GPT-4 results was 0.72 (range: 0.48 [Fear] – 0.89 [Valence]). The average NPV, with the more lenient threshold, was 1.00 (range: 0.99 [Valence] – 1.00 [Happiness]). **Figure 8** (right panel) shows the emotion-specific PPVs. Most findings in the image experiment did not survive the conservative thresholding (p < 0.05, voxel-level FWE-corrected) either for human- or GPT-based results making the PPVs and NPVs for the conservative thresholding meaningless.

**Figure 8.**
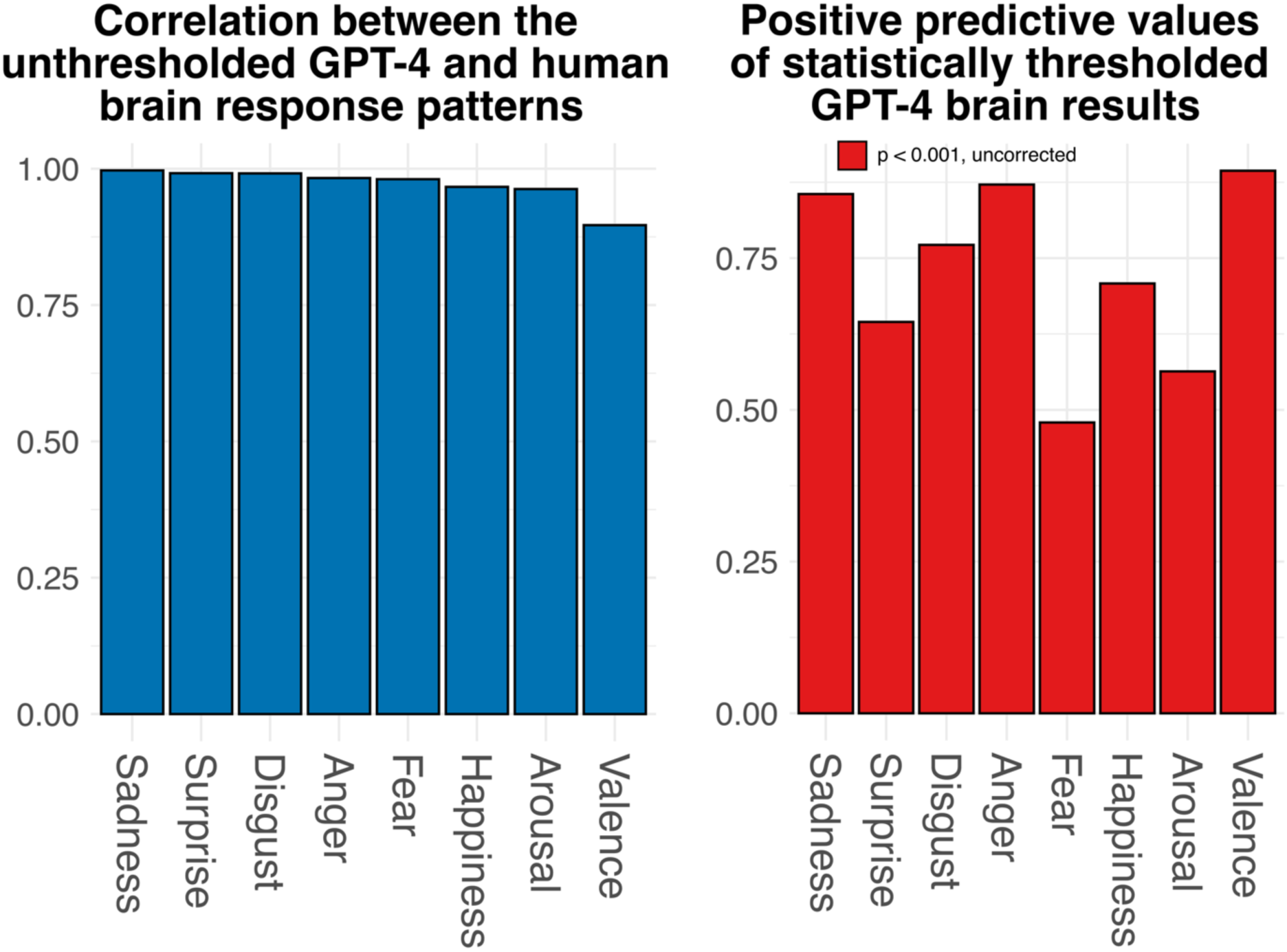
Similarity of the neural response patterns for the image fMRI experiment. The left graph shows the spatial correlation of the whole-brain response maps between GPT-4 and human results. The right graph displays the statistically thresholded positive predictive values (for positive association between BOLD signal and emotion) of the GPT-4 results.

### Neural organization of emotional processing based on GPT-4 evaluations

**Figure 9** shows cumulative brain activation patterns calculated as the sum over statistically thresholded response patterns for specific emotions (p < 0.001, uncorrected) to highlight the areas associated with emotional experiences. Consequently, the maps indicate how many emotions the BOLD signal in each brain region is associated with, indicating the brevity of tuning for different emotions. The cumulative maps for emotions based on GPT-4 ratings demonstrated a high degree of similarity to the cumulative map derived from human ratings in both video and image fMRI experiments (r_VD1_ = 0.95, r_ID_ = 0.85).

**Figure 9.**
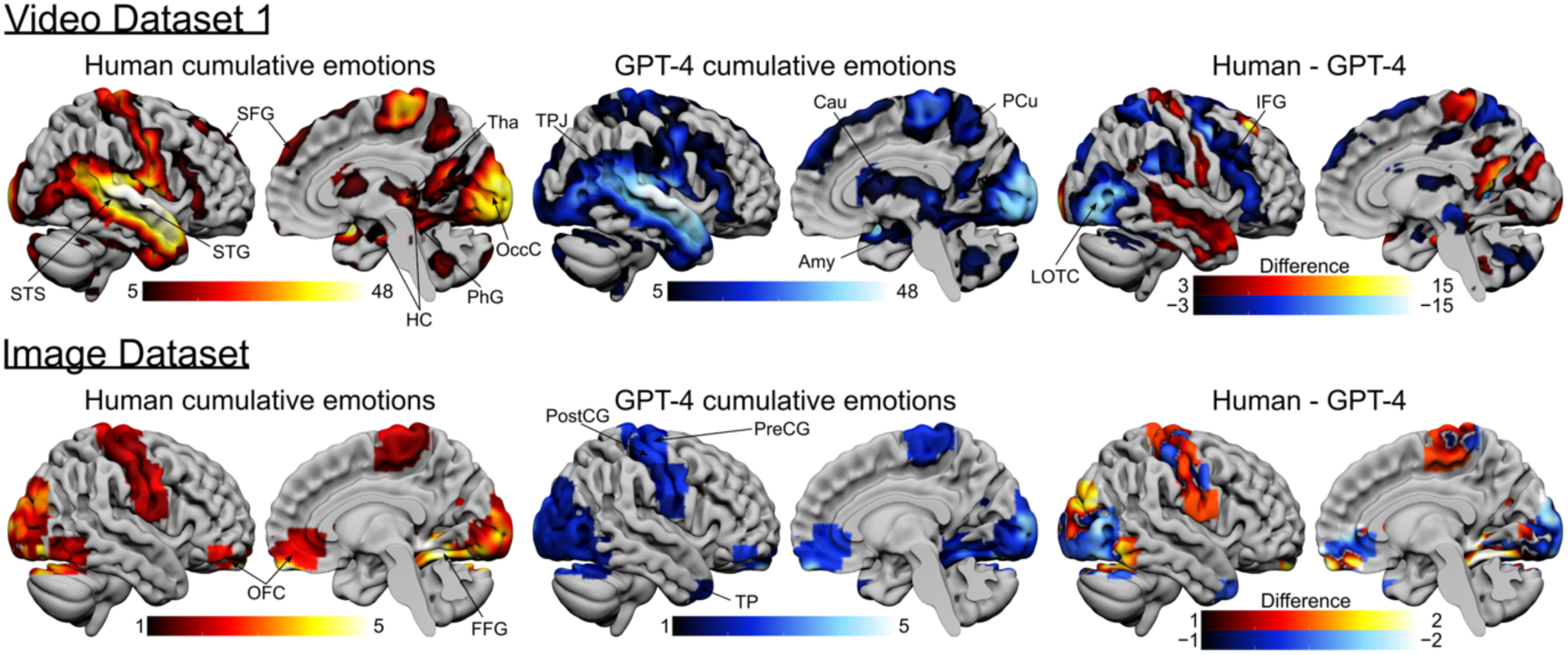
Organization of emotional circuits based on human and GPT-4 emotion evaluations. The surface maps show the number of emotions (out of all 48 emotions for videos and out of 8 for images) that were positively associated (p < 0.001, uncorrected) with the BOLD response. The top row shows the cumulative maps derived from the video fMRI experiment, while the bottom row shows results for the image fMRI experiment. The left column shows the cumulative maps based on human stimulus models, and the middle column shows the cumulative results for GPT-4 stimulus models. The right column shows the difference between the two cumulative maps, so that hot colors indicate areas where the human cumulative map shows more associations with emotions compared to the GPT-4 cumulative map. Amy: Amygdala, Cau: Caudate nucleus, FFG: Fusiform gyrus, HC: Hippocampus, IFG: Inferior Frontal gyrus, LOTC: Lateral occipitotemporal cortex, OccC: Occipital cortex, OFC: Orbitofrontal cortex, PhG: Parahippocampal gyrus, PCu: Precuneus, PostCG: Postcentral gyrus, PreCG: Precentral gyrus, SFG: Superior frontal gyrus, STG: Superior temporal gyrus, STS: Superior temporal sulcus, TP: Temporal pole, TPJ: Temporoparietal junction, Tha: Thalamus.

Cumulative maps of emotions in the video fMRI experiment highlight the commonly identified emotion and social perception networks, including superior temporal sulcus (STS), superior temporal gyrus (STG), lateral occipitotemporal cortex (LOTC), temporoparietal junction (TPJ), Fusiform gyrus (FFG), inferior frontal gyrus (IFG), caudate nucleus (Cau), thalamus (Tha), Amygdala (Amy), Parahippocampal gyrus (PhG), Hippocampus (HC) and medial superior frontal gyrus (SFG). Cumulative maps for the static image experiment highlighted the orbitofrontal cortex (OFC), Fusiform gyrus (FFG), Precentral gyrus (PreCG), and Postcenral gyrus (PostCG).

## Discussion

Our main finding was that GPT-4 can accurately predict the emotions that humans experience when viewing dynamic video clips or static images. These responses formed a two-cluster structure around pleasant and unpleasant emotions, and the emotion structures accord between humans and GPT-4 as well as across independent datasets. Finally, functional brain circuits associated with emotional processing can be mapped with GPT-4-derived emotion annotations, yielding results that are comparable to those obtained with traditional stimulation models using human evaluations. These results are based on actual visual input, not just textual descriptions of situations, and they were replicated using datasets whose emotion ratings have never been published, preventing “leakage” of the knowledge of the material to GPT-4. These results highlight the GPT-4’s capabilities to predict how humans feel when presented with a large variety of visual stimuli, paving the way to broad application potential in emotion research, cognitive neuroscience, and practical solutions.

This research extends our previous work, where we showed that GPT-4 can evaluate the presence of complex social information in dynamic social situations (Santavirta et al. 2025). The previous study established that GPT-4 can extract complex social information from visual stimuli, but it did not investigate whether GPT-4 can predict human emotional ratings to such stimuli. Altogether these results show that GPT-4, and most likely other flagship LLMs, can describe the social contents of situations and predict human’s emotional experiences for visual stimulation surprisingly well. The overall correlation of emotion ratings between GPT-4 and human was between 0.71 and 0.8 across datasets. Previous studies have already established that LLMs are able to recognize emotions from text descriptions. For example, LLMs can solve standard emotional intelligence tasks (Schlegel, Sommer, and Mortillaro 2025) and they exhibit elements of cognitive empathy (Sorin et al. 2024). Especially GPT-4 possesses significant capabilities in understanding situational emotions (Sabour et al. 2024; Tak and Gratch 2024; X. Wang et al. 2023; Tak, Gratch, and Scherer 2025). Our results go significantly beyond these data because in real-life emotions are mainly evoked by dynamic audiovisual information where emotional and social cues are often complex and context dependent. Such situations are difficult to describe accurately in text, and the present results extend these findings to more life-like conditions and natural emotional perception of visual scenes.

### Consistent ratings for experienced emotions between GPT-4 and humans

The GPT-4 emotion evaluations were computed as the average of ten independent evaluation rounds for the same stimuli to ensure the stability of the evaluations. Taking the average of repeated GPT-4 evaluations also increased the consistency between GPT-4 and human evaluations (**Figure SI-1**) similarly as we observed in the previous study for social information annotation (Santavirta et al. 2025). The increase in accuracy is most likely due to filtering out the randomness in the individual responses, or in other words, finding the most likely estimates produced by the internal probability distributions of the model.

With this approach the GPT-4 average ratings achieved high consistency with the human average annotations for 48 emotions (including 34 unipolar emotions and 14 affective dimensions) in two video datasets and for six basic emotions, valence and arousal for images (r_VD1_ = 0.71, r_VD2_ = 0.77, and r_VD2_ = 0.80, **Figure 2** & **Figure 5**). On the level of individual emotions, the median distance between the human and GPT-4 ratings were mostly small (median distance ≤ 1 indicating under 10% difference for 34 emotions in VD1, for 39 emotions in VD2, and for all six basic emotions in ID). Although the overall consistency with humans was high, GPT often gave lower ratings (typically 0 or 1 normalized units lower) than humans. This was most prominent for the cases when humans had evaluated their emotional responses to be mild (normalized rating ≤ 5, **Figure 2**). This difference was more clearly visible in VD1, which contained videos with dialogue and background music (VD2 only included silent videos). GPT-4 was fed with eight snapshots from a video clip along with the transcript, and therefore it did not have access to all visual and auditory information such as the tone of human voice or background music that may alter the emotional context of the situations (Cowen et al. 2019). Hence, humans had access to the most subtle cues that might have not been available for GPT-4, which could have resulted in GPT-4 responding that the emotional response was completely absent when humans reported slightly higher emotional responses. However, at higher values, this bias vanished (**Figure 2**), indicating that GPT-4 was more accurate in capturing strong emotional experiences. We anticipate that attempts to send even more information (especially auditory information) to GPT-4 might increase its accuracy even further.

Although the overall consistency with human was high for image evaluation as well, GPT-4 seemed to slightly overestimate rather than underestimate emotion ratings compared to humans (**Figure 5**, left graph). Although the NAPS BE dataset was originally curated to evoke basic emotions, the original human evaluations for this data indicated that the participants did not feel very strong emotions when watching these images. In contrast, our human participants reported also strong emotional responses (normalized rating ≥ 8) for the video stimuli suggesting that videos evoke strong emotions more effectively than images. The reason for GPT’s overestimation for images can thus be due to the lack of dynamic content needed to effectively evoke emotions in humans. However, this bias can also be an artefact stemming from differences in data collection and differences in the human rating samples across video and image datasets. While GPT-4 and humans had exactly same instructions to evaluate ratings on a scale 9-point Likert scale from "not at all" and 9 being "very much", the previously collected ratings for images used a visual analog scale (VAS) which could influence human evaluations differently than the Likert scale. Third possibility is that when humans watch images in quick succession, the emotional responses do not change very quickly, while GPT-4 as an artificial system is not temporally constrained in its responses.

### Consistency of GPT-4 emotion ratings compared to consistency between humans

Emotional experiences are subjective and variable across individuals despite being broadly consistent. Hence, we benchmarked the consistency of GPT-4 emotion ratings (correlation with the human average rating) by calculating intersubject consistencies (single participants rating correlation with others) and group-level consistencies (rating correlation between two groups of five participants) for the human sample. This analysis indicated that consistency of GPT-4 was, on average, higher than either of these (**Figure 4**). The consistency of GPT-4 exceeded the intersubject consistency for 96% (VD1) and 92% (VD2) of emotions and even exceeded the group-level consistency for 73% (VD1) and 46% (VD2) of emotions. The median consistency of GPT-4 was 0.65 (VD1) and 0.79 (VD2) while the human group-level consistencies were 0.62 and 0.79, respectively. These results indicate that GPT-4 can predict the population average emotional responses for video stimulation with higher accuracy than single human observers and the consistency is even higher than between two groups of five human observers.

### Consistent structural representation of emotion ratings between GPT-4 and humans

The structural similarity of emotion ratings between GPT-4 and human observers was high for all three datasets. The structural representation of emotion ratings was identified by calculating the correlation matrices of emotions from the original ratings. Correlation matrices were calculated separately for GPT-4 and human data and also for each dataset to allow cross-comparison (**Figure 6**). Within datasets the structural representations of emotions were similar between GPT-4 and human ratings (r_VD1_=0.84, r_VD2_=0.88, and r_ID_=0.97). Additionally, the correlation matrices were consistent across VD1 and VD2 indicating that the emotion representations form mainly a two-cluster solution that distinguishes pleasant emotions from unpleasant ones. All cross-correlations between the correlation matrices of the two datasets (VD1_human_ vs. VD2_human_, VD1_GPT_ vs. VD2_GPT_, VD1_human_ vs. VD2_GPT_, VD1_GPT_ vs. VD2_human_) were over 0.8. These results indicate that the emotional space across 48 emotions is stable with different video stimuli and between human and GPT-4 annotations.

### Neural circuits of emotional processing modeled with GPT-4 emotion predictions

To demonstrate the utility and reliability of GPT-4 emotion ratings in cognitive neuroscience, we modeled the neural representation of emotion circuits using retrospective fMRI datasets where emotions were elicited to 97 healthy participants with VD1 videos and ID images. The hemodynamic responses were modeled separately with GPT-4 and human ratings to allow comparative analysis of the results. Previous analyses based on ratings revealed high levels of consistency for self-reported feelings, and as expected, this consistency extended to the neural level.

Cumulative activation maps highlighted a broad, human-typical socioemotional processing network in the video stimulation dataset (**Figure 9**, top panel). The cumulative maps were remarkably similar when the data were modeled using GPT-4 versus human emotion ratings (r_VD1_=0.95 and r_ID_=0.85). This analysis highlighted superior temporal sulcus (STS), superior temporal gyrus (STG), lateral occipitotemporal cortex (LOTC), temporoparietal junction (TPJ), Fusiform gyrus (FFG), inferior frontal gyrus (IFG), caudate nucleus (Cau), thalamus (Tha), Amygdala (Amy), Parahippocampal gyrus (PhG), Hippocampus (HC) and medial superior frontal gyrus (SFG) as the main hubs for socioemotional processing for the VD1 social video stimulation. Cumulative maps for the static image experiment highlighted the orbitofrontal cortex (OFC), Fusiform gyrus (FFG), Precentral gyrus (PreCG), and Postcenral gyrus (PostCG). According to previous studies, these regions are central to emotion elicitation and response, emotion recognition, and especially processing of socioemotional cues, such as facial expressions, speech prosody, and body language (Lettieri et al. 2019; Saarimaki et al. 2016; Koide-Majima, Nakai, and Nishimoto 2020; Saarimäki et al. 2025).

When investigating the convergence at the level of individual emotions, the results were also highly consistent between humans and GPT-4. The average correlation of unthresholded beta-maps for each emotion was 0.91 for VD1 and 0.97 for ID. However, in VD1, slightly lower consistency (r < 0.80) was observed for emotion "awe", "obstruction", "disgust", and "effort" which may be due to the ambiguity or their context-related nature. With respect to the ID, the individual correlations for the dimensions of valence and arousal, and for the six emotions, were all above 0.80. The average positive predictive value (PPV) of the GPT-4-based results for VD1 was 0.80 for both conservative (FWE-corrected, p < 0.05) and lenient (uncorrected, p < 0.001) statistical thresholds and the average negative predictive value (NPV) was 0.97 (FWE-corrected) and 0.96 (uncorrected), respectively. These metrics were highly similar in the ID. This suggests that the GPT-4-based models can predict neural emotion circuits associated with specific emotions with high similarity to the gold-standard human-annotation based models.

### The effect of the chosen GPT model and content moderation

The reported GPT-4 annotations were collected using OpenAI’s GPT-4.1 (gpt-4.1-2025-04-14) model but we initially collected the evaluations with GPT-4 Turbo model (gpt-4-turbo-2024-04-09). A comparison of the results between these two models is presented in **Table SI-5**. The performance of GPT-4.1 was minimally better compared to the older model. The most significant difference between the models was not their accuracy in making assessments, but rather the increased content moderation in the newer model. While GPT-4 Turbo provided emotional assessments for nearly all stimuli, GPT-4.1 refused to evaluate a few videos (1-2 %), mainly including sexual content, as well as a few images (1%) containing blood, wounds, or nudity. This suggests that the GPT-4.1 model has tightened internal content filters deliberately limiting its use cases with certain material. Content that raises or addresses conflicting moral opinions is also likely excluded from the training data. According to the GPT-4 model documentation, the content filters include, for example, "sexual," "hate," "violence," and "harassment" filters (https://platform.openai.com/docs/api-reference/moderations/object). While these filters may promote the safe and child-friendly use of LLM models, they also limit their usability in studies examining basic human behavior such as sexuality and aggression. This restricts the range of material that can be evaluated and may introduce bias if certain emotional domains (e.g. sexuality) are overlooked systematically.

With closed source-models, such as GPT-4, it is currently not possible to turn oF content moderation. Open-source models could be useful tools for research when commercial content moderation significantly limits the use of closed source models. However, currently open-source models are inferior compared to GPT-4 in multiple benchmarks, and they require significant local computational resources. One major benefit for closed-source models is how easily they can be accessed without strong computational background or resources, making them available for wide audiences. For these reasons closed-source models like GPT-4 may strike a practical balance between performance and accessibility.

### Cost-efficiency

The potential economic impact of automated approach to emotion annotation is considerable. For example, in the current study, the total cost of collecting emotion ratings from human observers exceeded 3000 dollars in participant fees, requiring a total of 260 hours of the participants’ time (∼29 min/participant). In contrast, the collection of GPT-4 estimates for all datasets using the API was fast and costed roughly 100 dollars (around $0.02 per video query and < $0.01 per image query, **Table SI-5**), or just 3% of the cost of human data. Automated annotations are not only highly cost-efficient, but they also overcome issues such as subject compliance or vigilance. In the future, laborious and expensive human experiments could be more efficiently targeted to test the most promising hypotheses, which could be first generated in pilot studies or in preliminary analyses with LLMs, such as GPT-4, before being fully and specifically tested with human participants.

### Future directions and applications

The results provide a strong foundation for utilizing GPT-4 and other large language models for studying emotions and their neural bases. The neural results demonstrate the potential to automatically annotate complex psychological phenomena for dynamic stimuli and then reliably model functional neuroimaging data with these annotations. This opens new avenues for automated approaches in large-scale neuroimaging experiments. In multidimensional and laborious stimulus mapping experiments, human annotations could be, at least partially, replaced by GPT-4 estimates (Dillion et al. 2023). This approach would accelerate the mapping of representation spaces, increase statistical power, and significantly reduce costs compared to previous multidimensional projects (Huth et al. 2016; Koide-Majima, Nakai, and Nishimoto 2020; Tarhan and Konkle 2020; Santavirta et al. 2023). For example, automated solution would allow re-annotating retrospective large-scale movie fMRI datasets in unprecedented detail and temporal scale easily and cost-efficiently for advanced analyses. Although the focus of this paper was to utilize LLMs for research purposes, automatic and real-time evaluations of complex socioemotional information from videos would have significant application potential in diverse real-life applications in surveillance and security systems, patient monitoring, customer experience analysis, and social robotics development, for example.

Currently, GPT-4 cannot simultaneously process high-frame-rate video with its full audio stream. Future research should attempt to increase the given information, especially auditory input, when prompting video evaluations from LLMs. However, we anticipate that full multimodality for native video input to the LLMs is nearby. While the present study used GPT-4, several other LLMs are available, and new versions of GPT as well as new generations of LLMs will very certainly soon emerge. Our general approach and specific results may provide a rationale for testing both existing and upcoming models: a new interdisciplinary subfield - at the crossroads of psychology, neuroscience and artificial intelligence – could propose standard procedures to evaluate specific LLMs with respect to their capacity to predict the outcomes of human psychological processes.

### Limitations

Overall, the results supported the hypothesis that GPT-4 can indeed predict human emotional evaluations for video and image stimulation with accuracy that is higher than agreement between small groups of people. However, our human data consisted of ratings from ten participants for each stimulus. Collecting larger human sample might yield higher confidence in the overall population average and would also allow estimating the consistency between larger subgroups of humans, such as testing LLMs’ performance in predicting emotions in different cultures. However, data collection costs would also increase significantly as described above.

The responses produced by LLMs are also sensitive to how the prompts are formulated. Previous studies have demonstrated significant impact of prompt wording on LLM model results (L. Wang et al. 2024), while we have previously found that minor changes in the prompts do not significantly change how GPT-4 evaluate social information from videos (Santavirta et al. 2025). However, the same problem applies also to humans, who can interpret even simple instructions or adjectives such as “happy” in vastly different ways. Nevertheless, we stress that we used a consistent prompt that closely mimicked the instructions given to human observers while ensuring that GPT-4 produced its evaluations in a structured format.

Like other LLMs, GPT-4 is trained on large and non-disclosed datasets most likely collected from the internet, books, and other sources. This exposes the models to underlying cultural and societal biases. Consequently, LLMs’ responses could be biased towards perspectives of a specific populations, ignoring the experiences and interpretations of other groups (Demszky et al. 2023). However, research on these biases is inconclusive (Santurkar et al. 2023; Park, Schoenegger, and Zhu 2024; Almeida et al. 2024), and the rapid evolution of models makes it difficult to keep up with investigating the properties of any specific version. While the precise impact of these biases on our estimates is unknown, our study focused on predicting population average emotional responses rather than ideological attitudes. Hence, it is reasonable to assume that the impact of potential biases on emotion assessments is minimal. Additionally, the human ratings included participants from over 30 nationalities ensuring diversity in the reference data. The human data for video datasets was collected for this study and they have not been made public before publications, which ensures that they have not being included in the GPT-4 training.

## Conclusions

Our study provides the first empirical demonstration that a large language model - GPT-4 - can accurately predict the intensity of 48 distinct emotions that humans feel when viewing a large array of videos and static images. GPT-4’s emotional ratings were found to be highly consistent with human average ratings. This consistency was higher than agreement between two groups of five human participants for most emotions. The structural representations of emotions were consistent between GTP-4 and human ratings, and also across two independent video datasets. Modelling the neural circuits for emotional processing in fMRI datasets with GPT-4 derived stimulation models predicted similar neural response patterns compared to traditional models based on human annotations. This opens significant possibilities in cognitive and affective neuroscience research where laborious multimodal human annotation can be complemented or sometimes even replaced with LLM-based annotations, allowing also large-scale reanalysis of existing datasets with novel stimulus models. Altogether our results show that multimodal language models provide a valuable tool for investigating the self-reported human affective experience and its neural basis.

## Data and code availability

The anonymized GPT-4 and human rating data and the collection and analysis scripts are available in the project’s GitHub repository (https://github.com/Lauri301/GPT-4-predicts-human-emotions). According to Finnish legislation, the original (even anonymized) neuroimaging data used in the experiment cannot be released for public use. The voxelwise (unthresholded) result maps from fMRI analyses can be requested from the authors. The stimulus movie clips of VD1 can be made available for researchers upon request, but copyrights preclude public redistribution of them. Short descriptions of each movie clip in VD1 can be found in the supplementary materials (**Table SI-1**). VD2 and ID are standardized datasets available from the original authors.

## CRediT authorship contribution statement

Severi Santavirta: Conceptualization, Methodology, Software, Validation, Formal analysis, Investigation, Resources, Data curation, Writing – original draft, Writing – review & editing, Visualization, Project administration. Lauri Suominen: Methodology, Software, Validation, Formal analysis, Investigation, Resources, Data curation, Writing – original draft, Writing – review & editing, Visualization. Yuhang Wu: Conceptualization, Methodology, Software, Validation, Resources, Writing – review & editing. David Sander: Conceptualization, Writing – review & editing. Lauri Nummenmaa: Conceptualization, Methodology, Resources, Writing – original draft, Writing – review & editing, Visualization, Supervision, Project administration.

## Acknowledgements

The study was supported by Turku University Foundation and Alfred Kordelin Foundation grants to SS and Finnish Governmental Research Funding for Turku University Hospital and for the Western Finland collaborative area to SS.

