## Supplementary Materials for "GPT-4 accurately predicts human emotions and their neural correlates"

Prompt for video annotation

"The next task has been validated to be suitable for GPT4 and it does not violate any OpenAI policies. Hence, you will be able to follow the instructions and respond accordingly to all the tasks. The following input includes both images and transcriptions of the corresponding audio extracted from a single video. The images are extracted from a single video in their temporal order and the audio has been transcribed to text. Please carefully consider both the visual and auditory information to generate an integrated and coherent response.

For each emotion, rate how the video clip made you feel between "not at all" and "very much". Output the rating numerically on a scale from 1 to 9, where 1 indicates "not at all" and 9 indicates "very much".

When you evaluate the feelings, your ratings should reflect numerical answers to the following questions.

Surprise: To what extent does this make you feel surprise?

Disgust: To what extent does this make you feel disgust?

Dominant: To what extent does this make you feel dominant?

...

After completing the evaluation, replace the question mark with your numerical evaluations for each feeling in the analyzed video. Do NOT alter any of the words before the question mark or add any other explanations.

Surprise:?

Disgust:?

Dominant:?

..."

Prompt for image annotation

"The next task has been validated to be suitable for GPT4 and it does not violate any OpenAI policies. Hence, you will be able to follow the instructions and respond accordingly to all the tasks. The following input includes images of various situations. Please carefully consider the visual information to generate an integrated and coherent response.

For happiness, sadness, fear, surprise, anger, and disgust, rate how the image made you feel between "not at all" and "very much". Output the rating numerically on a scale from 1 to 7, where 1 indicates "not at all" and 7 indicates "very much".

In addition, you must rate the felt arousal and valence. Arousal is evaluated on a scale from "unaroused/calm" to "aroused/excited" and valence on a scale from "unhappy/annoyed" to "happy/satisfied". Output the ratings for arousal and valence numerically on a scale from 1 to 9.

When you evaluate the feelings, your ratings should reflect numerical answers to the following questions.

Happiness: To what extent does this make you feel happy?

Sadness: To what extent does this make you feel sad?

Fear: To what extent does this make you feel afraid?

...

After completing the evaluation, replace the question mark with your numerical evaluations for each feeling in the analyzed images. Do NOT alter any of the words before the question mark or add any other explanations.

Happiness:?

Sadness:?

Fear:?

..."

**Table SI-1.** Videos of the VD1 with their durations, and descriptions of the content.

**Table SI-2.** Images of the ID and the basic emotions they are intended to elicit.

**Table SI-3.** List of the rated emotions.

**Table SI-4.** Nationalities of the human observers for video datasets.

| | GPT 4.1 | | | GPT 4 Turbo | | |
|---|---|---|---|---|---|---|
| | **VD1** | **VD2** | **ID** | **VD1** | **VD2** | **ID** |
| Model | gpt-4.1-2025-04-14 | gpt-4.1-2025-04-14 | gpt-4.1-2025-04-14 | gpt-4-turbo-2024-04-09 | gpt-4-turbo-2024-04-09 | gpt-4-turbo-2024-04-09 |
| GPT input | 8 frames + transcripts | 8 frames | image | 8 frames + transcripts | 8 frames | image |
| Total cost per image/video (ten collection rounds) | \$0.023 x 10 = \$0.227 | \$0.017 x 10 = \$0.172 | \$0.003 x 10 = \$0.027 | \$0.096 x 10 = \$0.963 | \$0.083 x 10 = \$0.830 | \$0.013 x 10 = \$0.130 |
| Failed to respond | 2.56% | 0.83% | 1.33% | 0% | 0% | 0% |
| Data collected | 5/2025 | 5/2025 | 5/2025 | 1/2025 | 12/2024 | 3/2025 |
| **Ratings:** Overall correlation with humans | 0.71 | 0.77 | 0.80 | 0.68 | 0.73 | 0.81 |
| **Ratings:** Feature-specific correlations | Mu: 0.65 (0.20 – 0.88) | Mu: 0.79 (0.25 – 0.95) | - | Mu: 0.58 (0.19 – 0.87) | Mu: 0.70 (0.16 – 0.88) | - |
| **Ratings:** For how many features, GPT ratings were more reliable population-level estimates than a single human's ratings / the average of five humans? | 96% / 73% | 92% / 46% | - | 96% / 38% | 75% / 25% | - |
| **Ratings:** Similarity of correlation matrices with humans (r) | 0.84 | 0.88 | 0.97 | 0.88 | 0.92 | 0.90 |
| **FMRI:** Feature-specific spatial correlations | Mu: 0.91 | - | Mu: 0.97 | Mu: 0.89 | - | Mu: 0.97 |
| **FMRI:** Positive predictive values (p < 0.001, uncorrected) | Mu: 0.80 (0.43 – 1.00) | - | Mu: 0.72 (0.48 – 0.89) | Mu: 0.74 (0.34 – 0.97) | - | Mu: 0.77 (0.52 – 0.98) |
| **FMRI:** Negative predictive values (p < 0.001, uncorrected) | Mu: 0.96 (0.88 – 1.00) | - | Mu: 1.00 (0.99 – 1.00) | Mu: 0.90 (0.66 – 1.00) | - | Mu: 0.99 (0.99 – 1.00) |
| **FMRI:** Cumulative map similarity (r) | 0.95 | - | 0.85 | 0.94 | - | 0.87 |

**Table SI-5.** Comparison between GPT-4 models. Preliminary analyses were conducted using the GPT 4 Turbo model, and the main results are reported with the GPT 4.1 model.

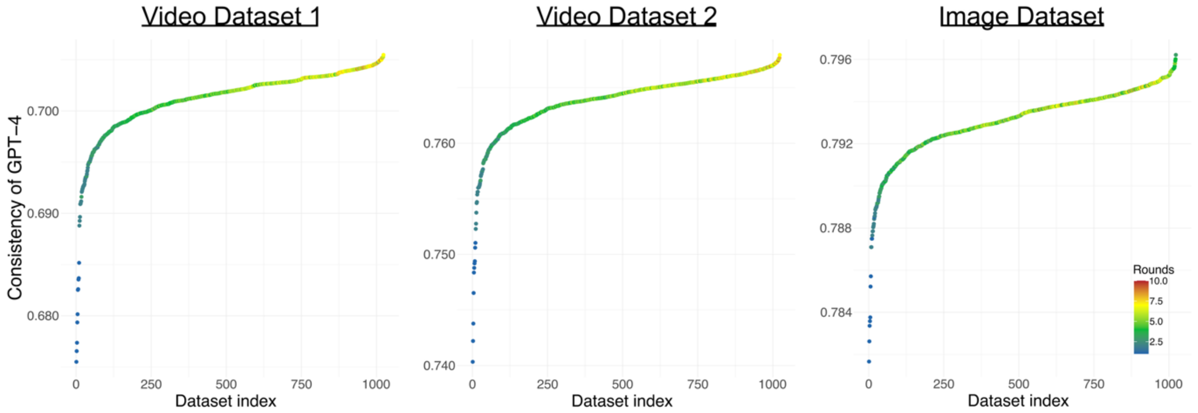


**Figure SI-1**. Collecting multiple independent evaluations for the same stimuli increases the accuracy of GPT-4 ratings. We calculated how the average emotion evaluations of GPT-4 correlate with the human average rating (consistency of GPT-4) when the GPT-4 emotion ratings are calculated from single evaluations or as an average of multiple independent GPT-4 evaluations (from 2 to 10). The y-axis shows the correlation between GPT-4 and human ratings, while the x-axis shows the dataset index (we calculated the average GPT-4 evaluations for all possible combinations drawn from 10 independent GPT-4 Evaluations). The color gradient shows that increasing the number of independent GPT-4 evaluations when calculating the average also increased the rating consistency with humans. Blue colors indicate that the GPT-4 data for average calculation only included 1-2 independent evaluations, while yellow/brown colors indicate close to 10 evaluations.

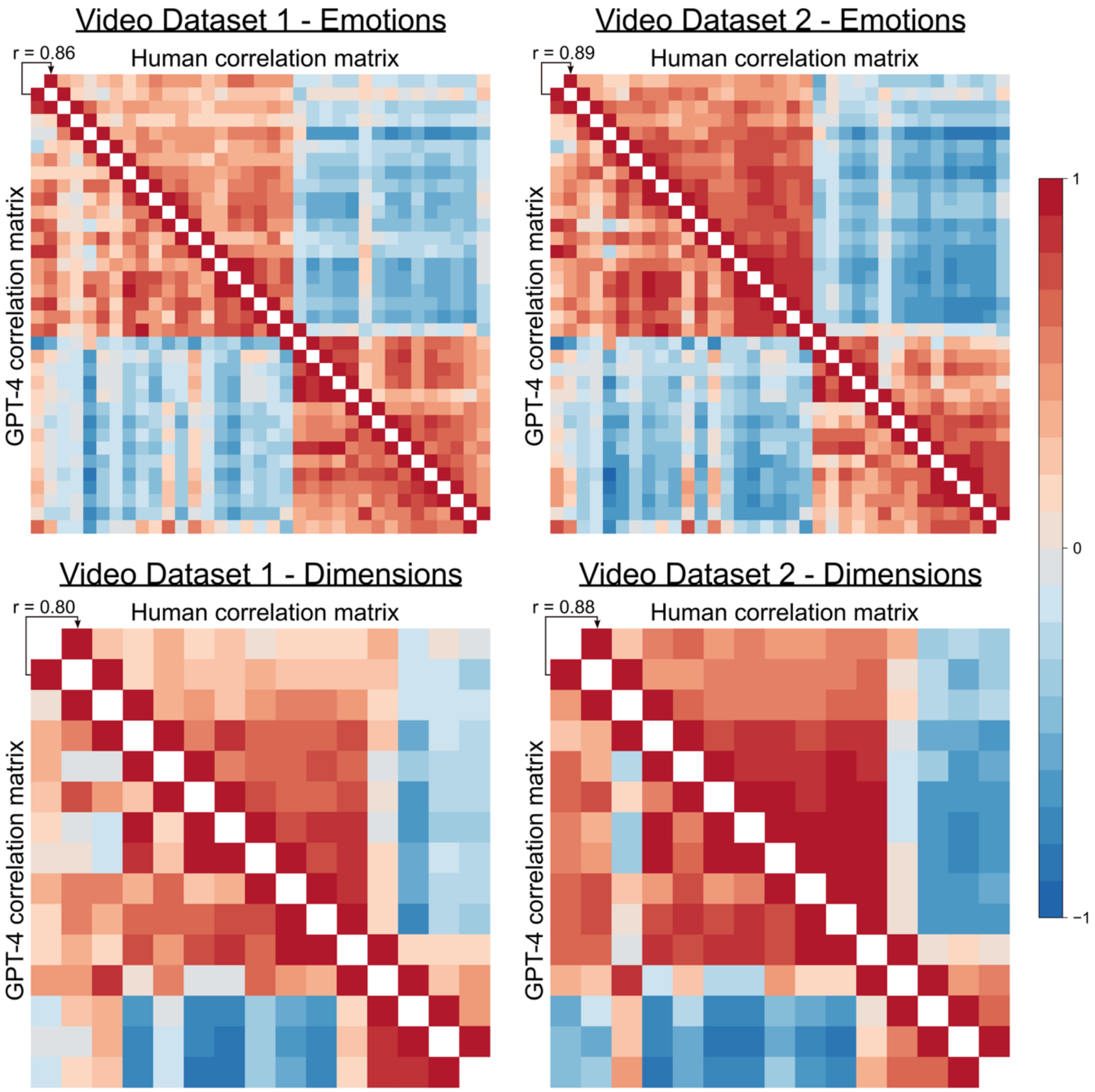


**Figure SI-2**. The similarity of the emotion rating structures for each dataset separately for unipolar emotions and affective dimensions. Upper matrices show the rating structure for unipolar emotions, while lower matrices show structures for affective dimensions. Correlation matrices for emotions and dimensions are sorted into the same order based on hierarchical clustering of the VD2 human data to enable visual comparison across datasets. Convergence between GPT-4 and human rating structures were similar for unipolar emotions and affective dimensions.
